# Invasion of epithelial cells are correlated with secretion of Biosurfactant via the type 3 secretion system (SST3) of *Shigella flexneri*

**DOI:** 10.1101/2020.04.02.021634

**Authors:** Duchel Jeanedvi Kinouani Kinavouidi, Christian Aimé Kayath, Etienne NGuimbi

## Abstract

Biosurfactants are amphipathic molecules produced by many microorganisms, usually bacteria, fungi and yeasts. They possess the property of reducing the tension of the membrane interfaces. No studies have been conducted on *Shigella* species showing their involvement of biosurfactant like molecules (BLM) in pathogenicity. This study aims to show that environmental and clinical strains of *Shigella* are able to produce BLM by emulsifying gasoline and diesel fuels. Our study has shown that BLM are secreted in the extracellular medium with EI24 ranging from 80 to 100%. The secretion is depending on the type III secretion system (T3SS). We did show that *S. flexneri*, *S. boydii* and *S. sonnei* are able to interact with hydrophobic areas with respectively 17.64%, 21.42% and 22.22% of hydrophobicity. 100 mM Benzoic and 1.5mg/mL Salycilic acids have been inhibited T3SS and this totally stops the BLM secretion. *Pseudomonas aeruginosa* which has T3SS is able to produce 100% of BLM in the presence or in the absence of both T3SS inhibitors. The secreted BLM is extractable with an organic solvent such as chloroform and could entirely be considered like lipopeptide or polypeptidic compound. By secreting BLM, *Shigella* is able to perform multicellular phenomena like “swarming” allowing to invade and disseminate inside epithelial cells.

## Introduction

The ingestion of pathogenic and virulent microorganisms generally affecting peoples in both developed and developing countries [1]. *Shigella* is one of the Gram-negative bacterium belonging to *Enterobacteriaceae* family and is causative agent of bacillary dysentery or shigellosis [2]. The genus *Shigella* was the major pathogen bacteria associated with dysentery with attributable fraction to 63,8%, but also the second most common pathogen associated with watery diarrhoea with attributable fraction to 12,9% in sub-Saharan Africa and south Asia. Children under 5 years are the most affected. More and more shigellosis is a pathology that both towards neglected diseases but 164300 of death per years have been notified all over the world in 2010. Most deaths occur in sub-Saharan Africa and in south Asia [3–6]. This is include Republic of Congo and surprisingly no epidemiological studies have been conducted in this field. The genus Shigella includes four species (*S. flexneri*, *S. sonnei*, *S. dysenteriae* and *S. boydii*) [7]. 10 bacteria of *S. dysenteriae* type 1 and 100 to 180 bacteria of *S. flexneri* or *S. sonnei* are enough to produce symptomatic infection [8].

*Shigella*’s pathogenicity is based on a virulence plasmid pWR100 in which the mxi-spa locus encodes the type three secretion system(T3SS) involved in effector production like IpaB, C and D (Translocator and Tip) to invade host cell [9–12]. A previous study in our laboratory that showed that *Shigella* sp. isolated from Brazzaville wastewater were able to emulsify hydrocarbon from gasoline and/or diesel fuel [13]. Sachin et al. found the same profile of hydrocrabon emulsification with *Shigella* strain [14]. According to amphipathic features, biosurfactants display a variety of surface activities, which explain their application in several fields related with emulsification, foaming, detergency, wetting, dispersion, pathogenicity and solubilisation of hydrophobic compounds[15, 16]. Biosurfactants are produced from microorganisms like *Pseudomonas aeruginosa*, *Bacillus subtilis*, *Candida albicans*, and *Acinetobacter calcoaceticus*. Rhamnolipids, sophorolipids, mannosylerithritol lipids, surfactin, and emulsan are well known and documented in terms of biotechnological applications [16–18].

*Shigella* pathogenicity mechanisms have been mostly studied using S. flexneri M90T as a model. In this study, we need to demonstrate that *Shigella* strains produce a biosurfactant in extracellular medium. Inner and outer membrane encompass numbers of secretion systems. In this way, this work aims to study the involvement of BLM via the type three secretion system (T3SS) pathways. In addition this work will assess the approvals that *Shigella* could use the BLM to promote the invasion and the dissemination inside epithelial cells.

## Material and methods

### Strains and Culture Conditions

Four (4) *Shigella* strains were kindly provided and collected from laboratory of Molecular Bacteriology (Free University of Brussels). This is include *S. flexneri* M90T, *S. flexneri spa40*-, *S. sonnei* and *S. boydii*. Three (3) pure culture strains were isolated from patients in Brazzaville University and Hospital Center (CHU-B) in 2018. These were provided by the Bacteriology Laboratory this hospital. Thirty (30) *Shigella* sp. strains were isolated in environmental wastewater of Brazzaville using decimal dillution in SS medium. *P. aeruginosa* [19] and E. coli Top10 were used as controls in this study. The strains were spread on the plates containing LB medium with Congo red with 100 μg/mL streptomycin for 24 hours at 37 ° C for wild type and 50 μg/mL for *spa40* mutant.

### Emulsification index (EI24) assay

An overnight of 5mL of bacterial culture have been done. The emulsification index (EI24) was calculated as an indicator for BLM as previously demonstrated [13]. The medium was adjusted to pH 7.2 and supplemented with gasoline or diesel fuel (1 mL for 300 mL of medium). The EI24 was investigated by adding fuels with LB medium in 1:1 ratio (v/v). The solution was vortexed for 5 min and incubated for 24 h at 37°C. The emulsification rate was calculated through the height of the emulsion layer. In addition, EI24 was determined for gasoline and diesel fuel hydrocarbons. All the experiments were performed in triplicates, EI24 = height of emulsion layer/total height of solution × 100.

### Bacterial swarming assays

Swarming was studied for all *Shigella* strain used in this study. Using plate assays containing 0.5% noble agar and LB medium with 0.5% dextrose. The mixture was sterilized at 121°C, during 15 min. After sterilization, the medium was supplemented with adequate antibiotics including streptomycin 100μg/mL for wild type and kanamycin 50 μg/mL for the Shigella flexneri spa40 mutant. Approximately 6 h after pouring the plates, bacteria were inoculated and spread by using a sterilized platinum wire with log-phase cells ([OD600] 0.6) grown in their respective media used for the swarming experiments. Swarming plates that were imaged only for their comparative end point swarming development were incubated at 30°C for 24 h prior to imaging [20].

### Bacterial Adhesion assay

The adhesion of bacteria to hydrophobic surface was evaluated according to the method described by Rosenberg [21]. The hydrophobicity was evaluated according to the following formula %H=A0-A/A0*100 with A_0_: OD before the mix; A: OD after vortexing of aqueous phase.

### Induction assay by using Congo Red

*Shigella* sp. have been cultivated in 5 mL of the final volume. 1 mL of overnight culture was fuged and 500 μL of sterile PBS and 10 μl of Congo red (10 mg / ml) have been gently added and mixed with the pellet by avoiding to break cells. Samples were incubated at 37 ° C with stirring. After 30 minutes of incubation, samples were centrifuged at 15.000 rpm for 15 minutes [11] at room temperature. Supernatants were gently recovered and mixture with gasoline or diesel fuels. The emulsification Index (EI24) have been determined.

### Extraction of Biosurfactant like molecule

Three methods have been used to extract biosurfactant.

#### HCL and Ethanol precipitation

This method was described by Vater [22]. An overnight culture has been centrifuged at 13,000 × g for 15 minutes. Once the supernatant was collected, HCl 1N and 90° ethanol were added to the supernatant. Precipitates have been generated by incubating samples at 4 ° C in overnight. Mixtures were fuged at 13000 g for 15 minutes to obtain granules. The granules obtained were tested with EI24 to evaluate the ability to emulsify the hydrocarbons.

#### Ammonium sulfate precipitation test

An overnight culture has been fuged at 13,000 rpm for 15 minutes to separate supernatant and pellet. Then 15 mL of supernatant were mixed with ammonium sulfate (80%) for 15 minutes. And finally this has been incubated with shaking overnight. The mixture has been fuged at 6000 rpm for 30 minutes at room temperature. Pelet has been hommogenized by using PBS buffer. The emulsification activity has been assessed.

#### Biosurfactant Extraction using Chloroform

The 24h culture was strictly centrifuged at 15,000 g for 15 minutes to avoid any residual bacteria. One volume of supernatant was added with an equal volume of chloroform (v/v). The mixture was strongly agitated by a vortex. After centrifugation at 6000 rpm for 10 min, the non-aqueous phase is recovered. The solvent was allowed to evaporate completely only without heating above 40°C. The residue is dissolved in a PBS buffer. The emulsification activity is tested by mixing with gasoline or diesel fuel in comparison with the supernatant at the start point. The emulsification Index (EI24) have been determined.

### Effect of Benzoic Acid and Salycilic acid on Biosurfactant secretion

Viability of *Shigella* strains have been first evaluated with different concentration of benzoic acid and salycilic acid. *S. flexneri* M90T was grown in Luria-Bertani broth (LB) in the presence of various concentrations of benzoic acid (50 mM, 100 mM, 250 mM and 500mM) and salycilic acid (1.5 mg/mL, 3 mg/mL, 6.25 mg/mL and 12.5 mg/mL). After that, all Shigella strain was grown in Luria-Bertani broth (LB) added with an adequate concentration of benzoic acid or salycilic acid at 37°C, during 24 hours. All supernatants were fuged and the secretion biosurfactants was assessed by using emulsification assay (EI24).

### Statistical analysis

GraphPad Prism 7 and Excel software were used for analysis. The data represent the arithmetical averages of at least three replicates. Data were expressed as mean ± SD and Student’s test was used to determine statistical differences between strains and p <0.05 was considered as significant. Principal Component Analysis (PCA) was used to investigate possible correlations between *Shigella* and emulsification index (EI24). Prior to ordination, percentage of emulsification activities data were transformed to better meet the assumptions of normality [23] using ln (x+1). All analysis was performed using CANOCO (Canonical Community Ordination, version 4.5) [24].

## Results

### Screening for biosurfactant production

In order to carry out our research, we first assess if *Shigella* strains are able to produce BLM in extracellular medium. As results Figure 1 showed that environmental strains and clinical strains are able to secrete BLM by showing emulsification percentages ranging between 15% to 100% (Figure 1A). *S. flex*neri spa40- was not able to produce BLM compare with *Pseudomonas aeruginosa* used as positive control. The way of strains to produce BLM is shown in figure 1B all strains are not represented (Figure 1B).

**Figure 1.**
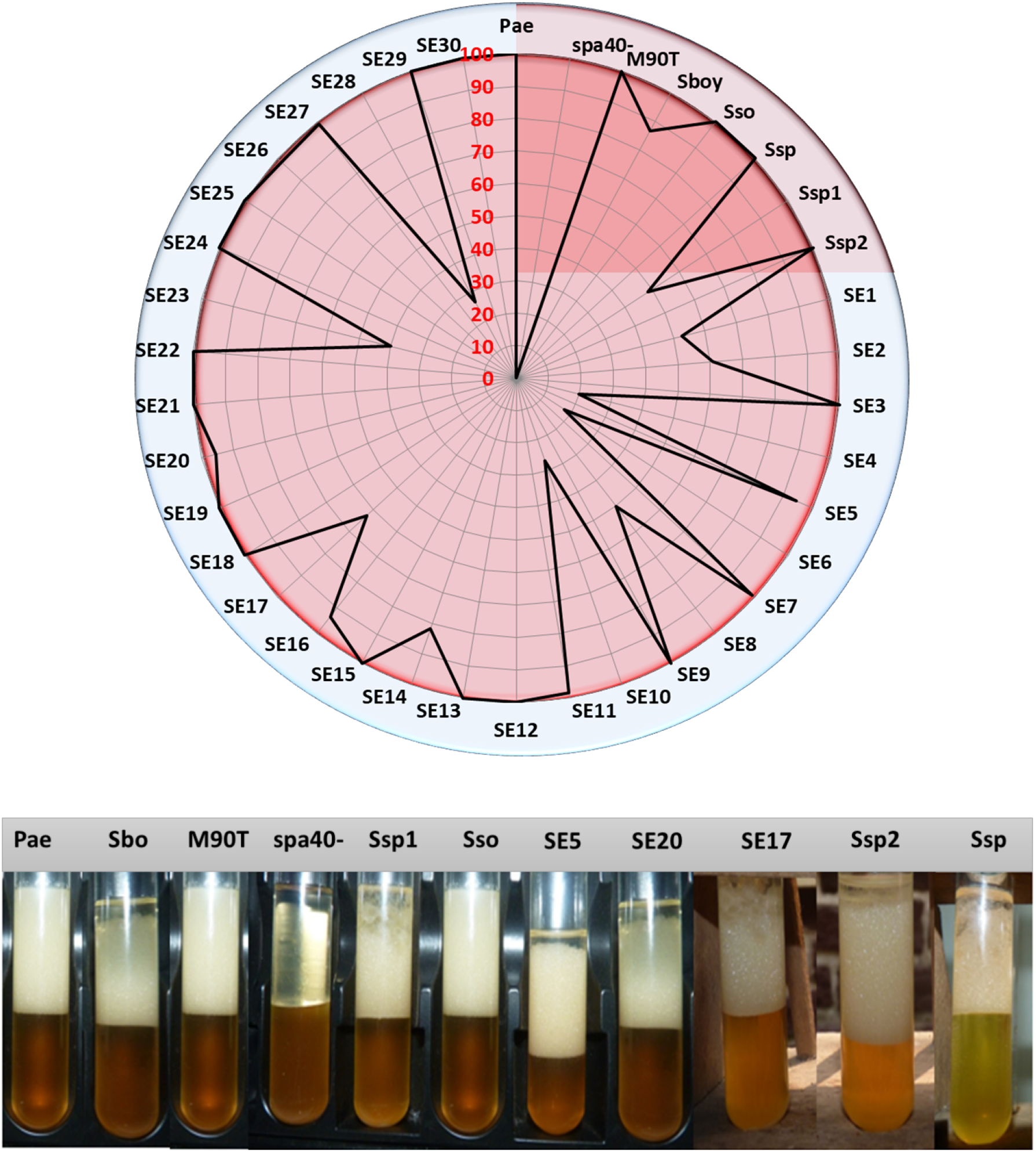
A. Emulsification index percentage of all *Shigella* strains used in this study after 24 hours. Pae: *P. aeruginosa* used as positive control; M90T: *Shigella flexneri* 5a strain M90T; *spa40*-: *S. flexneri* spa40 mutant; Sbo: *S. boydii*; *Sso*: *S. sonnei*; *Ssp, Ssp1, Ssp2*: *Shigella* sp. from clinical strains; SE1…30: *Shigella* sp. from environmental strains. B. Emulsification index appearance of some *Shigella* strains.

EI24 of Some strains are ranging from 80 to 100%. This is included *Shigella flexeneri* M90T, *Shigella boydii* (Sbo), *Shigella sonnei* (Sso), *Shigella* sp (Ssp), Ssp2, SE3, SE5, SE9, SE11, SE12, SE13, SE14, SE15, SE16, SE18, SE20, SE21, SE22, SEI24, SE25, SE2626, SE27, SE29 and SE30 (Figure 2). The positive control has been found in this rate. SE1, SE8, SE23 and Ssp1 are ranging between 40 and 60%. SE4, SE10 and SE28 are between 20 and 40%. SE17 and SE2 from 60 to 80% and Shigella flexneri spa40 mutant (spa40-) is not able to produce biosurfactant and SE6 is about 17% ranging from 0 to 20% (Figure 2).

**Figure 2:**
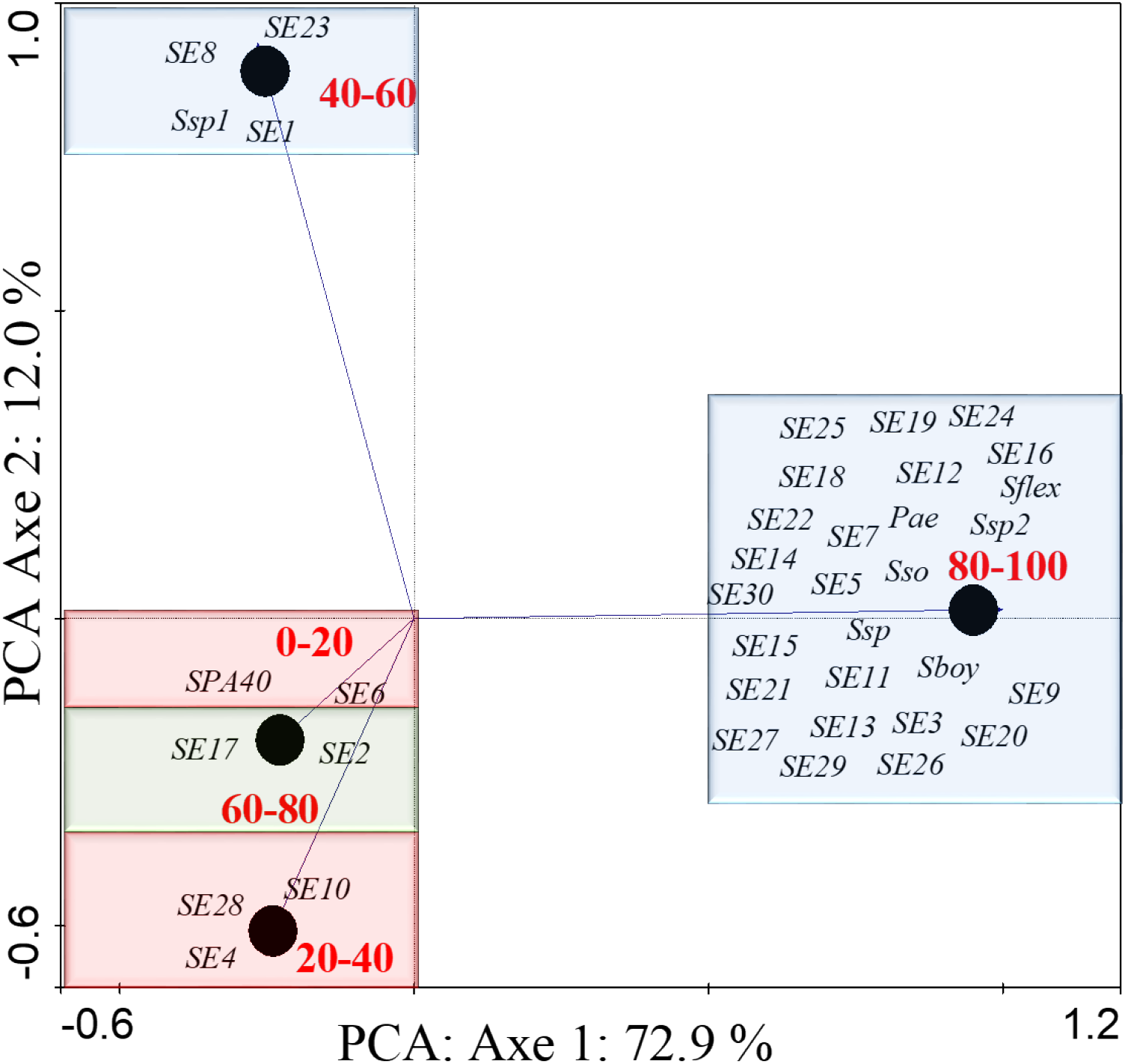
PCA of *Shigella* strains basing on emulsification index (EI24). Pae: P. aeruginosa used as positive control; S. flex: *S. flexneri* M90T; S. flex-: *S. flexneri spa40* mutant ; Sboy: *S. boydii*; S. so: *S. sonnei*; Ssp 1, 2: Shigella sp clinical strain; S. E1…30: *Shigella* sp environmental strain.

### The ability of Shigella strain in swarming test

Swarming is induced by the production of BLM. In order to demonstrate how *Shigella* can disseminate in epithelial cell, we first investigated if all *Shigella* strains used in this study were able to swarm by using (0.5%) LB medium + 0.5% dextrose. As result *S. flexneri* M90T, *S. sonnei* and *S. boydii* were able to spread and swarm. *spa40-* was not able to swarm (Figure 3). Some examples of the swarming profile of some *Shigella* strain after 24 hours are illustrated. We found that *S. flexneri spa40*- cannot swarm and *S. sonnei* have a particular swarming profile than other *Shigella* strain used in this study.

**Figure 3 :**
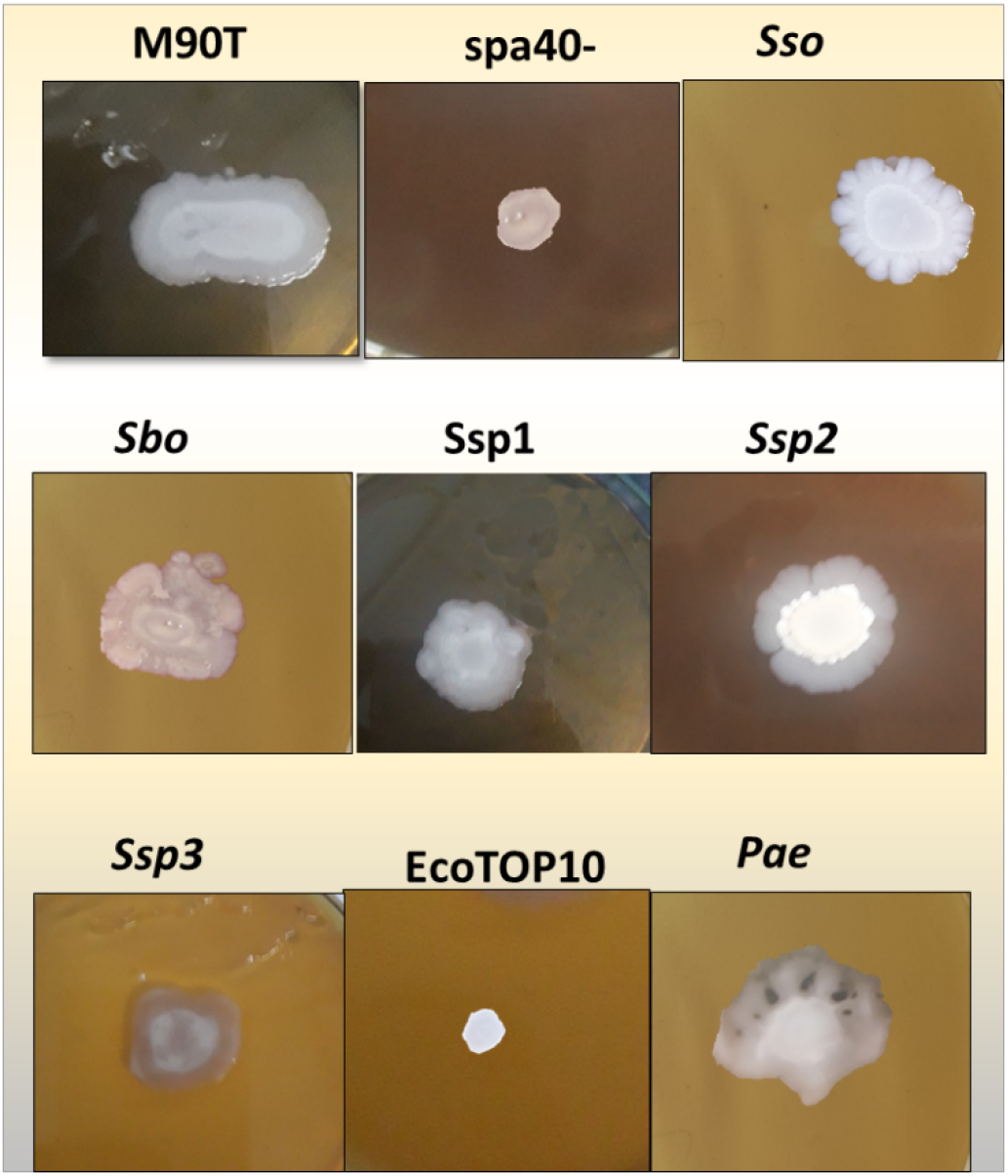
Swarming profile of *Shigella strains.* M90T*: Shigella flexneri* 5a strain M90T. Sbo: *S. boydii*) ; Sso: *S. sonnei* ; *spa40-*: *S. flexneri* 5a *spa40-* ; Ssp1, 2, 3: *Shigella* sp. Pae: P. aeruginosa used as positive control and E. coli-Top10 used as negative control.

### Bacterial Adhesion To Hydrocarbon (BATH)

To highlight the production of BLM by *Shigella* strains to induce interaction with hydrophobic areas, we performed analysis by evaluation of the ability to interact with hydrocarbon. The Figure 4 shows the bacteria adhesion profile of some *Shigella* strains. As results only *S. flexneri* Spa40 mutant does not interact with hydrocarbon area (Figure 4A). *S. flexneri*, *S. boydii* and *S. sonnei* are positive with BATH techniques including a percentage of hydrophobicity of 17.64%, 21.42% and 22.22% respectively (Figure 4B).

**Figure 4:**
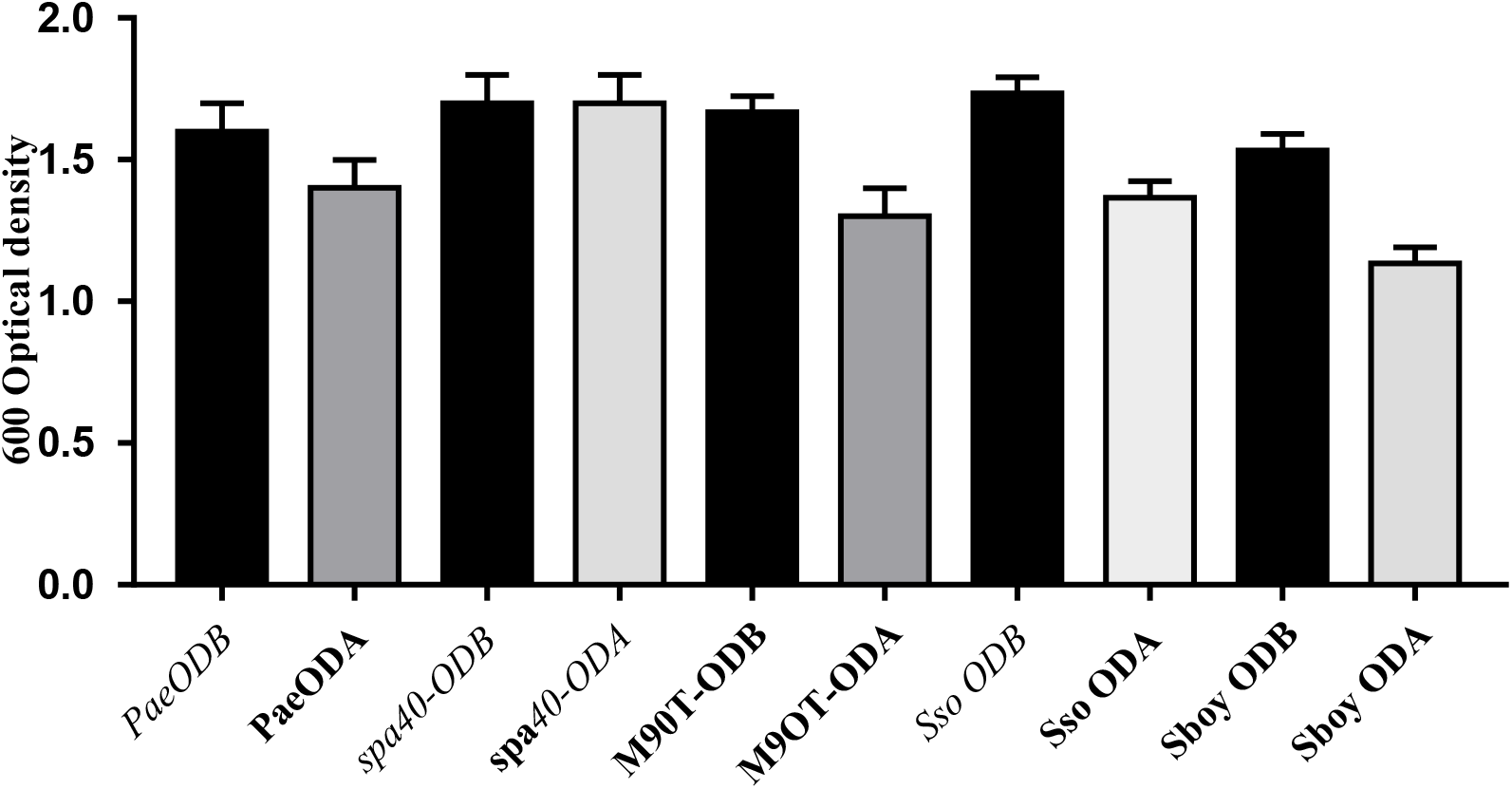
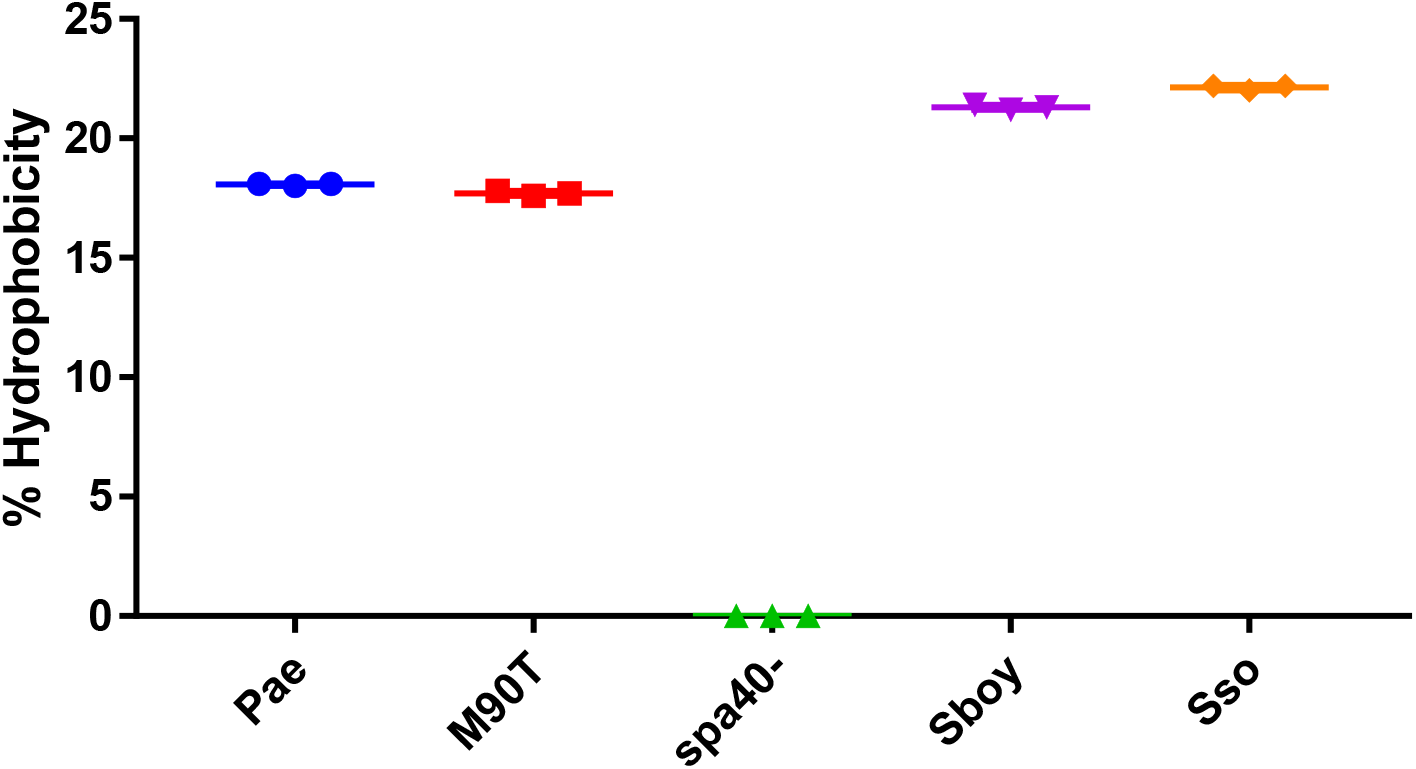
A. *Shigella*’s adhesion to hydrocarbon phase of some strains used in this study. ODB: optical density before vortexing. ODA: optical density after vortexing. M90T: *Shigella flexneri* 5a strain *M90T*. *Sbo:* S. boydii*) ; Sso: S. sonnei ;* spa40- : S. flexneri *5a* spa40-. Pae: *P. aeruginosa* used as positive control. B. Percentage hydrophobicity of *Shigella* strains.

### Screening of biosurfactant secreted by *Shigella* sp

To highlight the nature of the BLM secreted by *Shigella* strains used in this study. Cultures of *Shigella* strains whose supernatant emulsified hydrocarbons (gazoline or diesel fuel), have been used to identify the type of Biosurfactant like molecules. Precipitation on hydrochloric acid, ammonium sulphate and ethanol have been done. All strains showed a precipitate at the bottom of the tube. The emulsification index after precipitation has been carried on (EI24). Only precipitate profile performed from the *S. flexneri spa40-* supernatant did not emulsify the gas oil and/or diesel fuel. *S*. *flexneri, S*. *sonnei, S*. boydii and three Shigella sp. have 100 % of EI24 (**Figure 5**).

**Figure 5:**
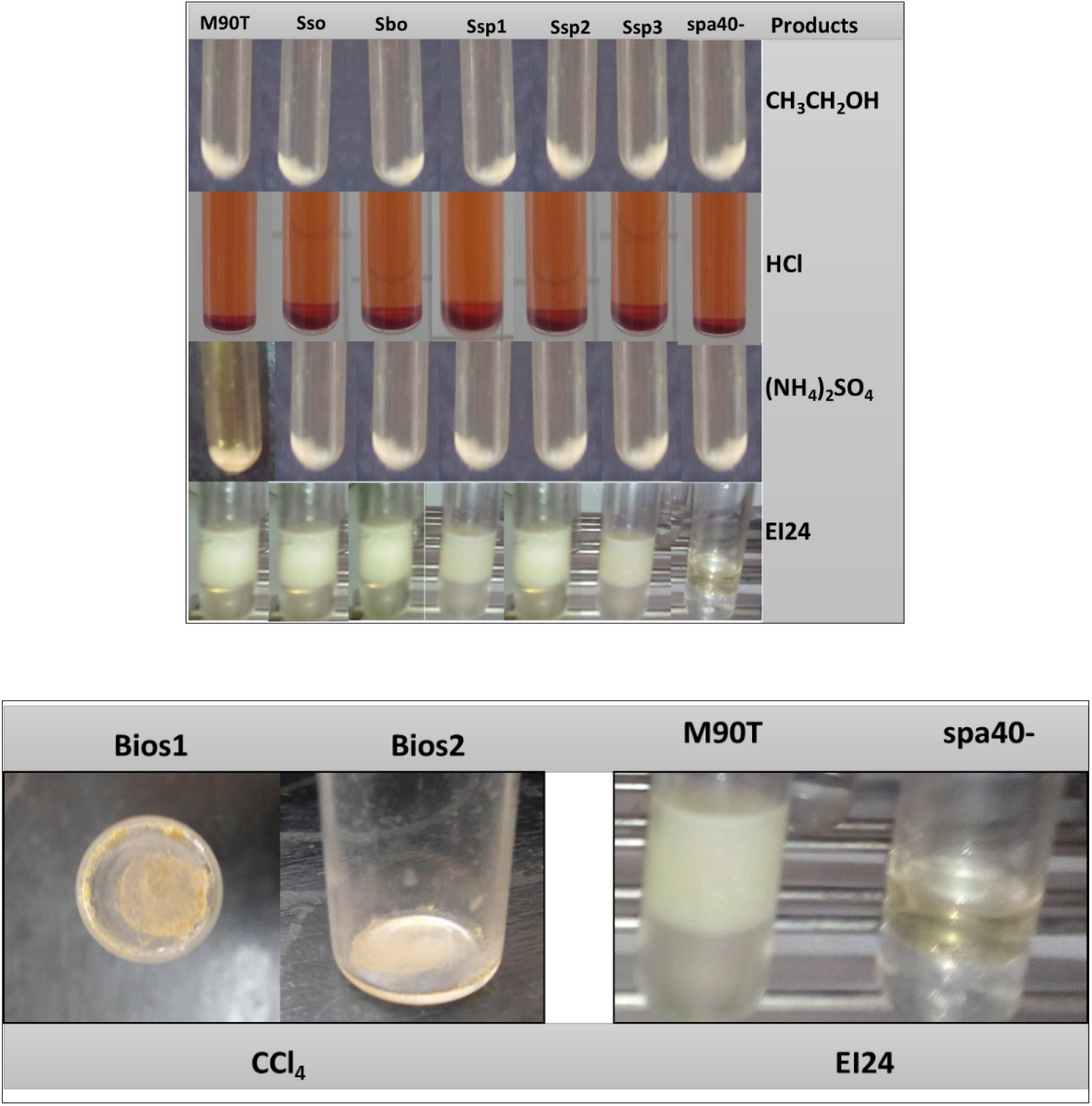
BLM purified from *Shigella Species*. TOP: Profile obtained after precipitation with ethanol (CH3CH2OH), Hydrochlorid acid (HCl) and ammonium sulfate ((NH4)2SO4). EI24: emulsification index for all strains (*S. flexneri* M90T, *S. sonnei*, S. *boydii*, *Shigella* sp.: Ssp1, 2, and 3. Down: left: residues obtained after evaporation of Chloroform (CCl4). Right: Emulsification index (EI24) for the extratable biosurfactant like molecule. Bios1 and Bios2: Biosurfactant like molecule residues.

Strains with known hydrocarbon emulsification ability were selected from an organic solvent like chloroform using biosurfactant extraction assay. Biosurfactant could be extracted after evaporation of chloroform at 40 ° C from *S. flexneri* M90T. Nothing was obtained from *spa40*- The extract after evaporation, suspended in PBS, was able to emulsify gasoline or gas oil with 100 % of EI24.

### The BLM is secreted by Type Three Secretion System (T3SS)

Clinical strains including *S. flexneri* M90T, *S. sonnei*, *S. boydii*, three *Shigella* sp. and 30 environmental strains including SE1 to SE30 were cultivated to induce the secretion of effector on Congo red induction. As results *Shigella* species have been found to secrete BLM on Congo red induction conditions with EI24 ranging from 80 to 100%. The mutant *S. flexneri* spa40- did not emulsify the gasoline and /or diesel fuel in the presence of Congo induction with 0% of EI24 (Figure 6A). Emulsification after Congo red type 3 secretion system of *Shigella* strains appearance are illustrated in Figure 6 down.

**Figure 6.**
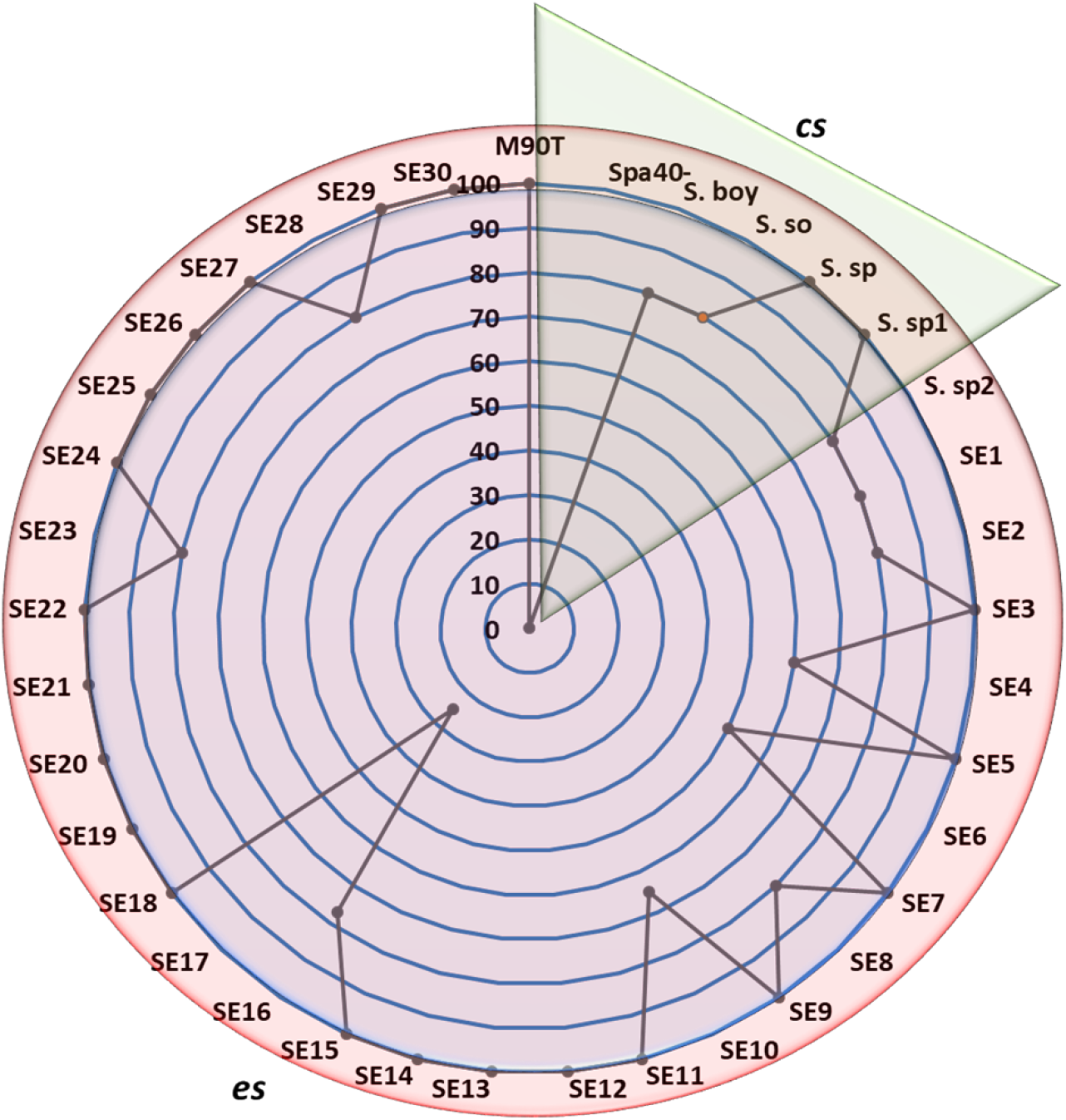
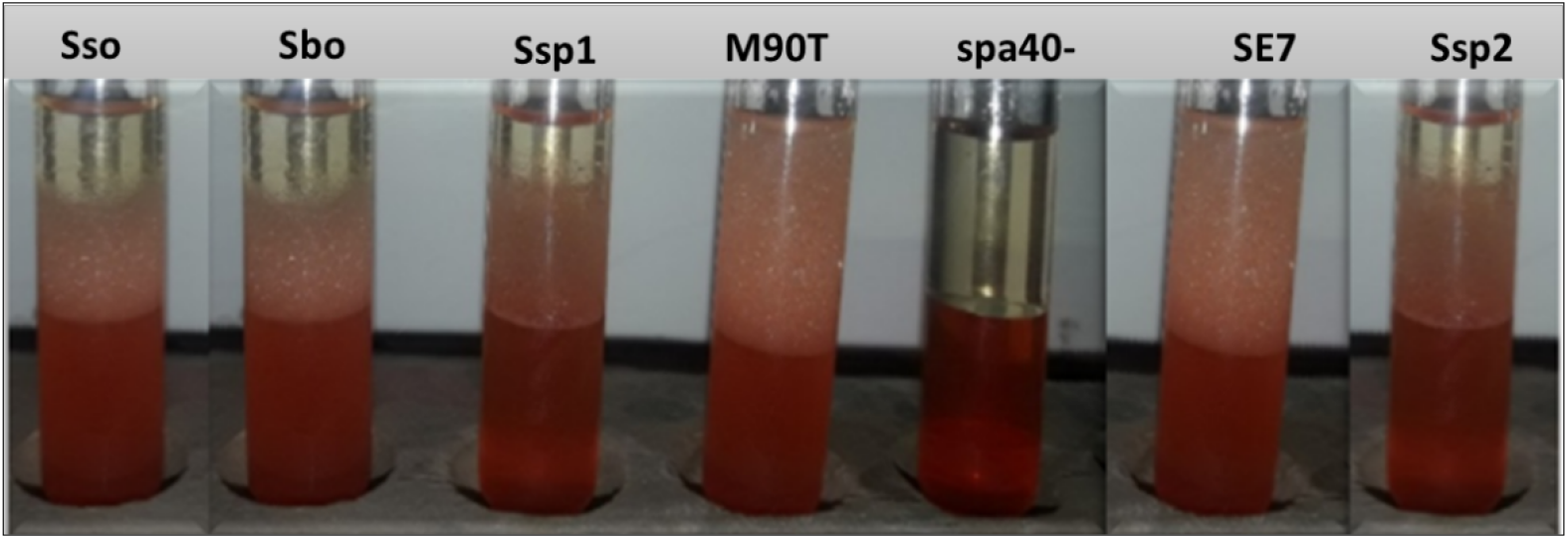
Emulsification index after Congo red induction. UP. Pae: *P. aeruginosa* used as positive control; M90T: *Shigella flexneri* 5a strain M90T; *spa40*-: *S. flexneri spa40* mutant; Sbo: *S. boydii*; *Sso*: *S. sonnei*; *Ssp, Ssp1, Ssp2*: *Shigella* sp. from clinical strains; SE1…30: *Shigella* sp. from environmental strains. Down. Emulsification index appearance of some *Shigella* strains.

### Effect of Benzoic Acid and Salycilic acid on Biosurfactant production

All *Shigella* strains were grown in Luria-Bertani broth (LB) in the presence of random concentrations of benzoic acid and salycilic acid (**Figure 7**). We examined growth at the various concentration of benzoic acid including 50 mM, 100 mM, 250 mM and 500 mM. As far as salycilic acid is concerned 1.5 mg/mL, 3 mg/mL, 6.25 mg/mL and 12.5 mg/mL were randomly selected. All *Shigella* strain grew normally within the physiological range of benzoic acid as determined by CFU per milliliter, but growth was significantly interesting at 100 mM benzoic acid and 1.5 mg/mL for salycilic acid (Figure 7).

**Figure 7.**
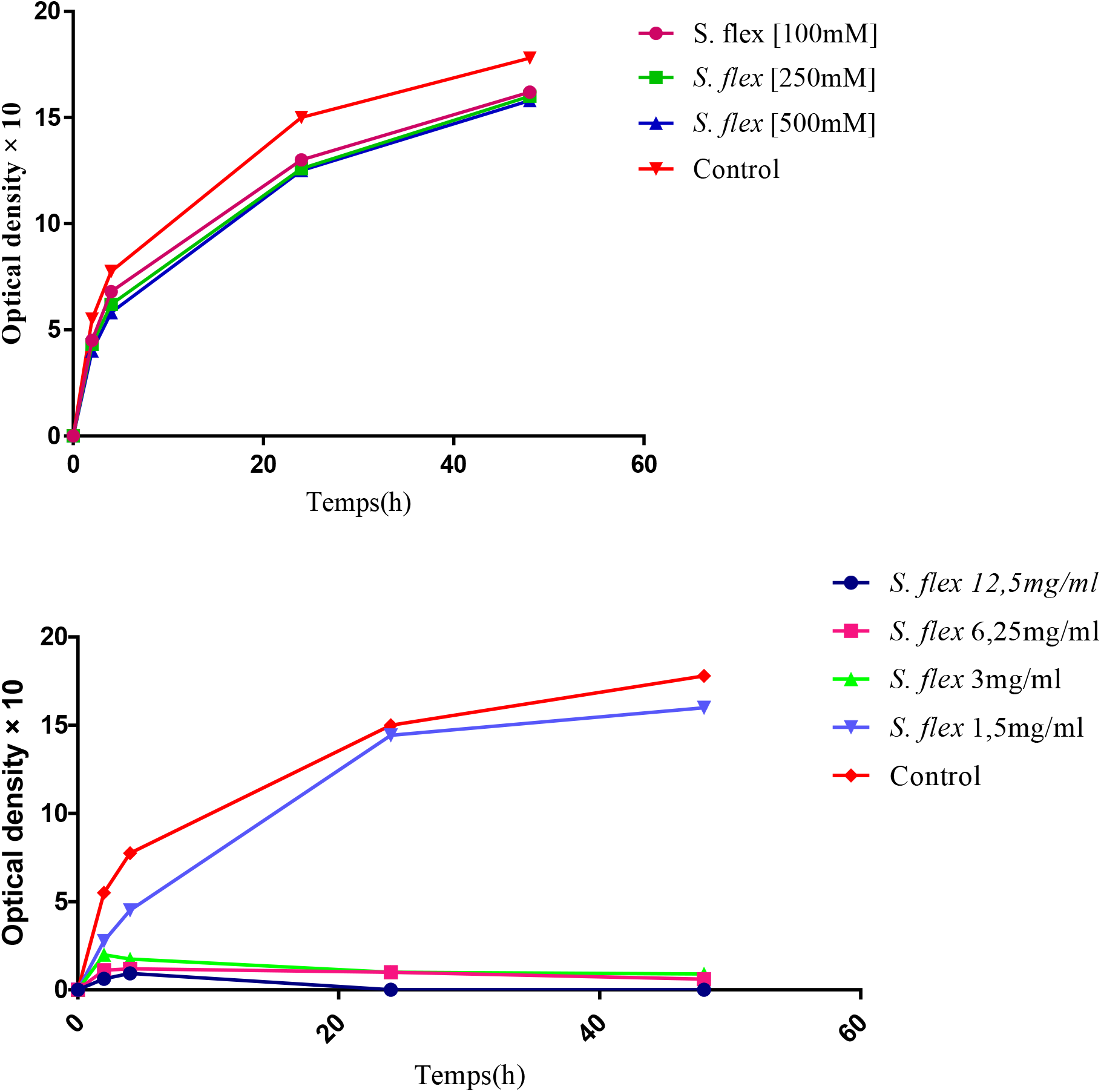
Growth curve analysis of all *Shigella* flexneri M90T used in this study on the presence of benzoic acid (TOP) and salycilic acid (Down).

To highlight the role of T3SS on the secretion of BLM we assessed the effect of benzoic and salycilic acids to inhibit the biosurfactant production. Bacteria were previously incubating with100 mM benzoic acid (BA) and 1.5 mg/ml salycilic acid (SA), we showed that *S. flexneri* M90T, *S*. *sonnei, S*. *boydii* and SE5 were not able to produce BLM with an emulsification index 0% EI24. This is easily showed that all *Shigella* strains do not emulsify anymore gasoline or diesel fuel with benzoic acid or salycilic acid (Figure 8 up). Strains are able to emulsify gasoline or diesel fuel without benzoic acid or salycilic acid (Figure 8 down). The appearance are illustrated in Figure 8 down. *P. aeruginosa* has been used as positive control since the T3SS is widely conserved in the most gram negative bacteria, surprisingly *P. aeruginosa* was able to produce 100% BLM in the presence or in the absence of BA and SA. It is worthy noticed that *spa40*- was not able to produce BLM neither in the presence nor in the absence as up mentionned (Figure 8).

**Figure 8:**
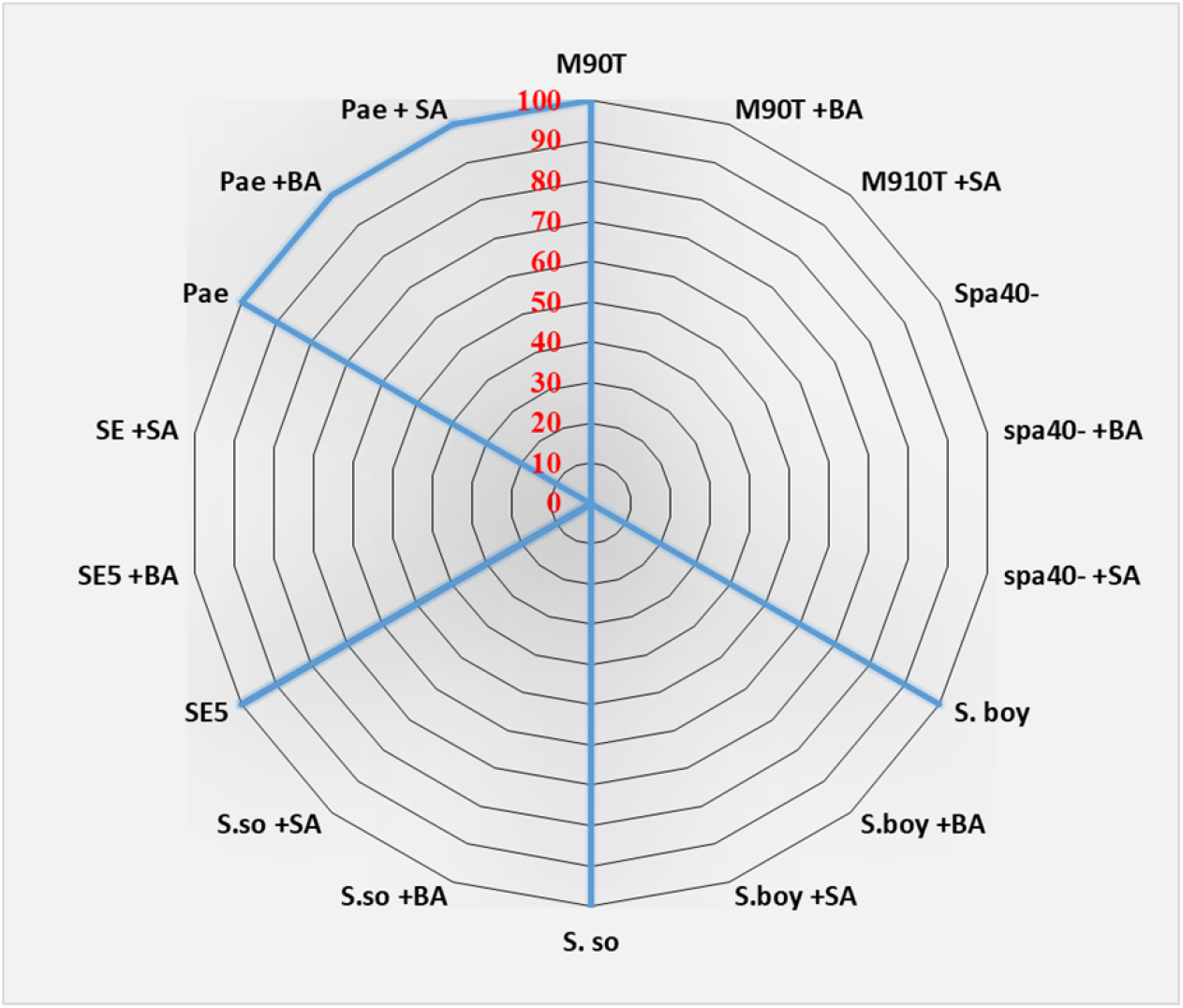
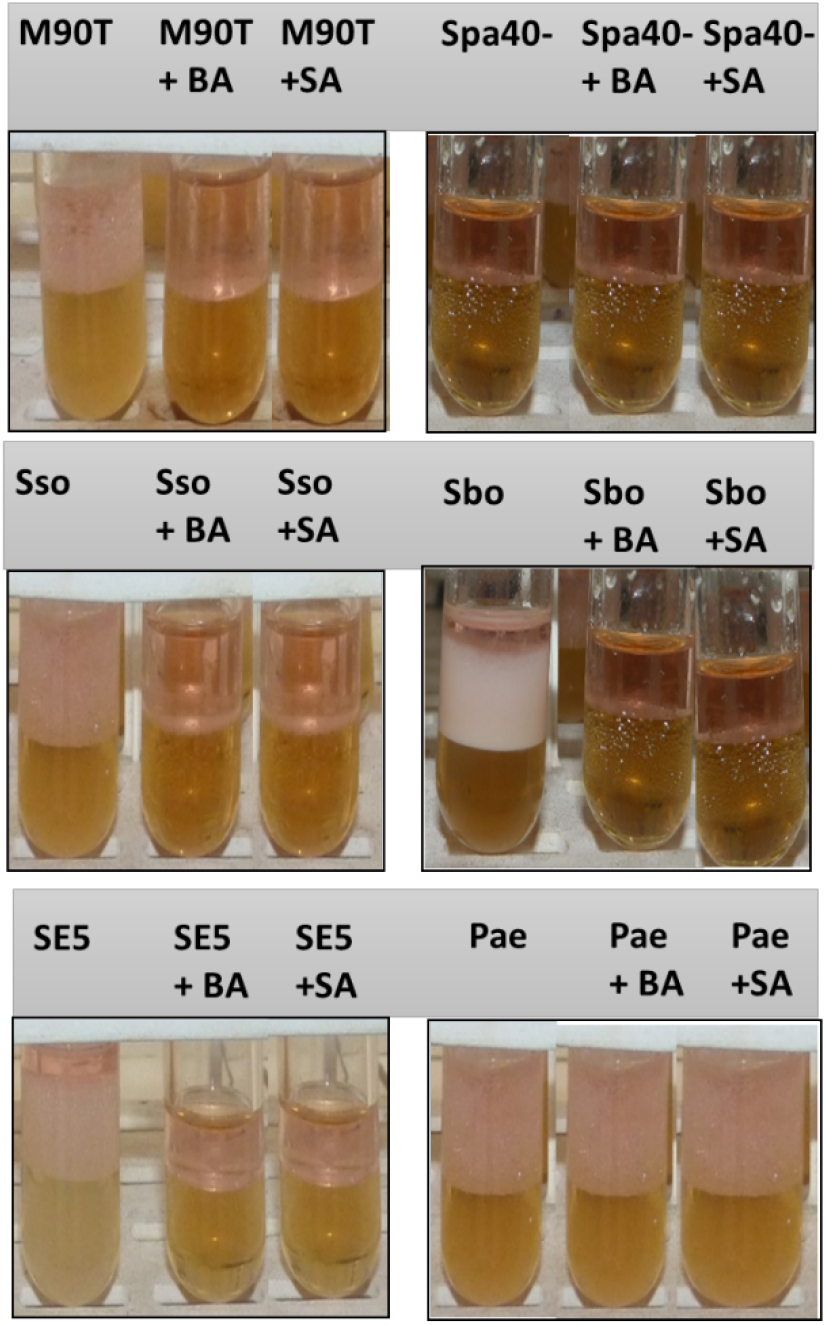
(a). Gasoline emulsifying activity of some *Shigella* strains used in this study with and without benzoic acid (BA) and salycilic acid (SA). M90T: *S. flexneri* strain M90T. spa40-: *S. flexneri spa40* mutant, *Sso* : *S. sonnei*; *Sbo*: *S. boydii*; *S.E*: *Shigella* sp (environmental strain) and Pae: *P. aeruginosa* (b). Gasoline emulsifying activity appearances of some *Shigella* strain tested with and without benzoic acid or Salycilic acid.

## Discussion

This work was conducted with the prime aim to contribute to the understanding of the *S. flexneri* 5a M90T epithelial cell invasion mechanisms. *Shigella* strains had been collected from the environmental areas, hospital or laboratory. All strains had the ability to produce BLM during growth in extracellular medium, and the production is strictly depending on T3SS pathway. This result shows very clearly that these molecules are secreted in the extracellular medium as described by Usman and al. [25]. *spa40* mutant which has no T3SS cannot secrete BLM. Several studies have been demonstrated the role of T3SS in the secretion of numbers of effector proteins involved in invasion and dissemination [26–33].

The emulsification index is a direct method for demonstrating the ability of strains to produce biosurfactants or not [13]. Those molecules have been known to form emulsions between two immiscible liquids [34, 35]. Experiments carried out from the acellular supernatant showed that *S. flexneri* 5a M90T as well as *S. boydii*, *S. sonnei* and other *Shigella* sp. are able to emulsifying gasoline and diesel fuels with EI24 ranging from 80 to 100%. Gram negative bacteria are well documented to overcome this phenomenon. This is include *P. aeruginosa* [34, 35], *Salmonella enteridis* [36], *Acinetobacter* sp. [37], and *Serratia Marcescens* [38]. Gram positive bacteria are known as professional one in the production of BLM. The spore-forming bacteria like *B. subtilis, B. lichenifornis* and *Lysinibacillus louembei* have been widely used to produce BLM [39–41].

Biosurfactants are native of several multicellular phenomena such as swarming described in several bacterial species [42–45]. By using specific culture media we have shown that all strain of *Shigella* genus were positive in swarming assay. The swarming phenomenon promote the ability for biosurfactants production. Since this phenomenon is associated with either antibiotic resistance, virulence and biofilm formation in *Proteus mirabilis*, *Salmonella enterica* serovar Typhimurium and *Serratia* [46–48]. This idea reinforces the fact that *Shigella* sp. could also use biosurfactants in its pathogenicity. No genes have been identified to be directly involved in BLM biosynthesis. In this work we found that *ygaG* is a chromosomal gene of *S. flexneri* M90T. YgaG which is the product of this gene shares 90 % of identity with LuxS involving in quorum sensing and biofilm formation [49, 50]. RhlA, RhlB and RhlR proteins are known to promote the rhamnolipid secretion [51]. The secretion of biosurfactant are correlated with quorum sensing [52].

Pathogenicity in genus *Shigella* is determined by T3SS that has the ability to secrete myriad of effector proteins into the target cell [11, 53]. In the absence of cellular contact, the secretory apparatus is not functional [54], however some proteins are secreted in leak condition. Cell contact is mimicking by the use of Congo red [55]. Under the Congo red induction condition, all *Shigella* strains emulsified gasoline and diesel fuels while the *S. flexneri* 5a M90T spa40 mutant did not emulsify anymore. The mutant *S. flexneri spa40*- has not T3SS [26]. The *S. flexneri spa40*- in non-inducible condition [56] or in a Congo red induction condition, does not produce BLM. In addition by blocking T3SS using benzoic and salycilic acids compounds, we have demonstrated that BLM could not be secreted in extracellular medium. This is confirm that BLM is secreted via T3SS. *P. aeruginosa* could secrete in the presence or in the absence of inhibitors. This allows us to postulate that rhamnolipid molecule could use another pathway. An efflux mechanism is the top in *P. aeruginosa* [25, 57]. This inhibition assay with benzoic acid and Salycilic acids showed a perfect correlation between the secretion of the BLM synthesized by *Shigella* and the inactivation of the type III secretion apparatus.

Regarding the BLM characteristics, precipitation assay like hydrochloric acid, ammonium sulfate and ethanol allowed to postulate that the secreted BLM could have a lipopeptide or peptide features. Only peptide or lipopeptide biosurfactants can precipitate at very low pH or with ammonium sulphate [22, 58]. In proteomics studies, the sequential precipitation of ammonium sulfate proteins allows the proteins to be separated by “salting-in” or “salting-out” effect [59], which necessarily leads to the formation of protein aggregates and therefore to their precipitation. The BLM precipitate was able to emulsify gasoline and diesel fuel. Biosurfactants like rhamnolipid, surfactin, emulsan are extractable by organic solvents [35, 60]. In addition our study showed that the biosurfactant excreted by *Shigella sp.* is extractable with chloroform with higher efficiency and stability at 40 ° C.

BML are known to play several vital roles especially in microbe’s adhesion, bioavailability, desorption and defense strategy. The most important role of microbial BLM is well reviewed for adhesion of the interfaces cells-cells interactions [61]. *P. aeruginosa* is a best example of cell surface hydrophobicity that justifies by the presence of cell-bound rhamnolipid [62]. Our new finding showed that by secreting BLM, *Shigella* sp. can easily bind to the cell hydrophobic interfaces by interacting with lipid rafts [27, 63–65]. By binding on cell membrane, BLM allows to reduce the membrane tension and to help the translocon like IpaB-C [66, 67] and the tip component IpaD [29, 32] to be closed to the host membrane and automatically to be inserted inside cytoplasmic membrane.

Many mechanisms have been demonstrated how *S. flexneri* can disseminate inside epithelial cells [68, 69], helping to escape autophagy phenomenon [70] and to spread inside host cell [71] by using a specific domain of IcsB that interacts with cholesterol [27]. In this work we showed that *S. flexneri*, *S. boydii* and *S. sonnei* can spread using swarming phenomenon. No studies have been previously documented about this strategy. This is efficiently emphasized and amplified the idea that *Shigella* is able to use several mechanisms that help *Shigella* to spread from cell to cell by secreting BLM. We are investigating the secretion of BLM inside epithelial cells. Basing on our finding, we can make a proposal that *Shigella* can invade and disseminate inside epithelial cells by using BLM.

## Conclusion

In order to contribute to the understanding of the mechanism of invasion of epithelial cells by *Shigella sp.* we have first shown that all *Shigella* strain as well as clinical strain or environmental strain are able to secrete biosurfactant like molecules directly in the extracellular. Secondly, we have shown that the secretion of biosurfactants like molecule are depending on Type Three Secretion System (T3SS). Our study suggest that, the biosurfactant with lipopetide or peptide features, stable at 40°C, could play an outstanding role on *Shigella* pathogenicity mechanisms including bacteria-host cell interaction, cell metabolism and cell dissemination. This work open ways to the understanding of genes associated with a couple of component that are able to promote the biosynthesis, the regulation and the secretion of BLM. MALDI-TOF and HPLC will be interesting to be done in order to characterize BLM.

## Data Availability

The Excel and Prism7 sheet including the data used to support the findings of this study are available from the corresponding author upon request.

## Conflicts of Interest

The authors declare that the research was conducted in the absence of any intellectual commercial or financial relationships that could be construed as potential conflicts of interest.

## Acknowledgments

We are grateful to Prof. Eric Déziel (Centre Armand-Frappier Santé Biotechnologie), Prof. Anne Botteaux (Free University of Brussels) for their continuous encouragements and for helpful data analysis before publication.

